# Investigating RNA-RNA interactions through computational and biophysical analysis

**DOI:** 10.1101/2022.02.01.478553

**Authors:** Tyler Mrozowich, Sean M. Park, Maria Waldl, Amy Henrickson, Scott Tersteeg, Corey R. Nelson, Anneke Deklerk, Borries Demeler, Ivo L. Hofacker, Michael T. Wolfinger, Trushar R. Patel

## Abstract

Numerous viruses utilize essential long-range RNA-RNA genome interactions, specifically flaviviruses. Using Japanese encephalitis virus (JEV) as a model system, we computationally predicted and then biophysically validated and characterized its long-range RNA-RNA genomic interaction. Using multiple RNA computation assessment programs, we determine the primary RNA-RNA interacting site among JEV isolates and numerous related viruses. Following *in vitro* transcription of RNA, we provide, for the first time, characterization of an RNA-RNA interaction using multi-angle light scattering (SEC-MALS) and analytical ultra-centrifugation (AUC). Next, we report the first RNA-RNA interaction study quantified by microscale thermophoresis (MST), demonstrating that the 5’ and 3’ TR of JEV interact with nM affinity, which is significantly reduced when the conserved cyclization sequence is not present. Furthermore, we perform computational kinetic analyses validating the cyclization sequence as the primary driver of this RNA-RNA interaction. Finally, we examined the 3-dimensional structure of the interaction using small-angle X-ray scattering, revealing a flexible yet stable interaction. This pathway can be adapted and utilized to study various viral and human long-non-coding RNA-RNA interactions, and determine their binding affinities, a critical pharmacological property of designing potential therapeutics.

**Graphical Abstract:** 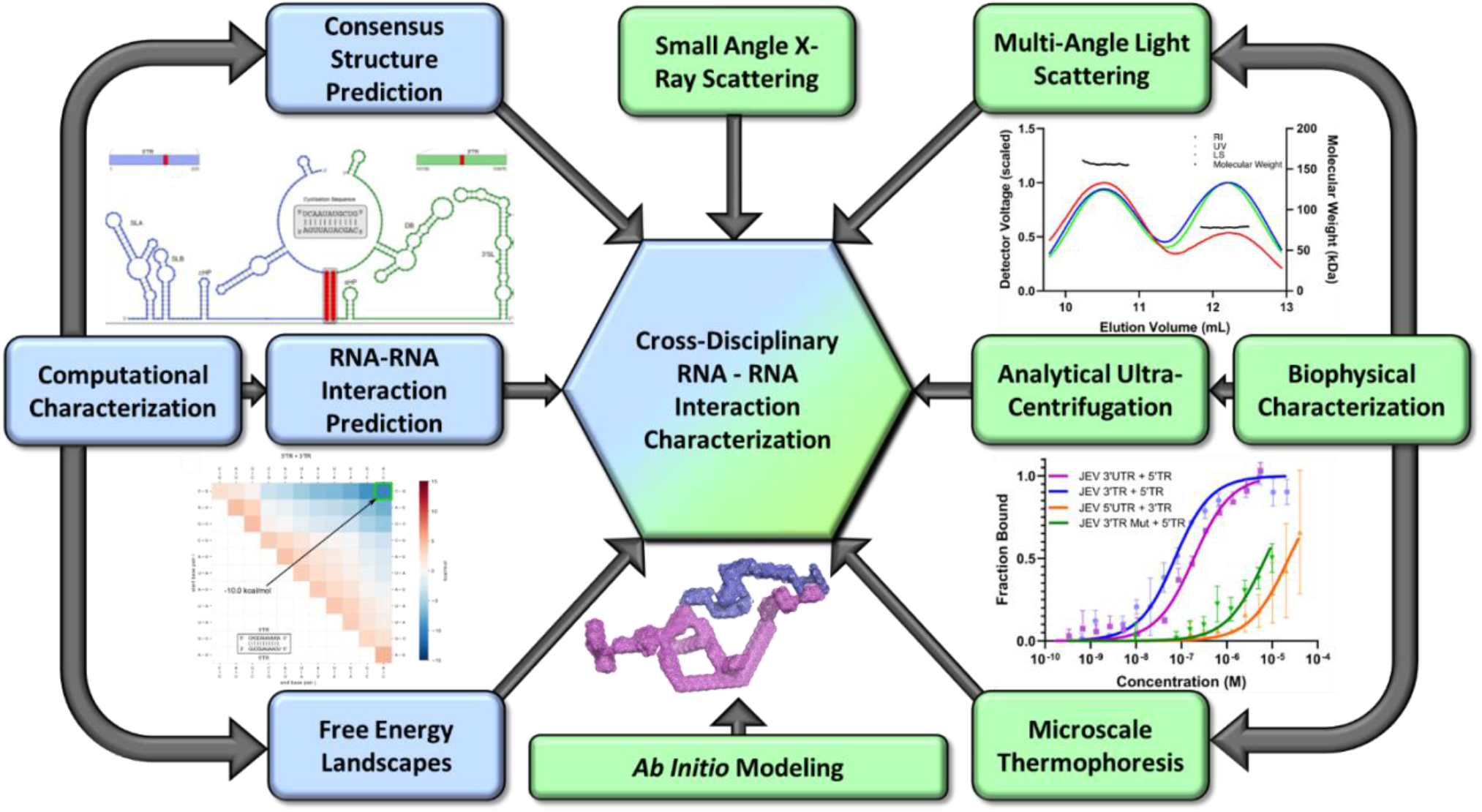

## Introduction

Computational and biophysical methods to obtain structure, properties, and dynamics of protein-ligand interactions are well developed as a foundational component of many pharmaceutical discovery pipelines. As a result, nearly all currently approved drugs target one of ~700 disease-related proteins, despite an increasing number of diseases attributed to the >98% of the human genome, which is non-coding (1). Amongst the limitations in targeting these elements is a lack of foundational techniques for characterizing RNA-RNA interactions in the same depth as protein-ligand interactions. The importance of RNA characterization has only gained traction as of late, highlighting an ever-increasing need for techniques capable of doing so (2–5). In this study, we describe the novel application of well-established biophysical modalities with computational studies for extensive characterization of RNA-RNA interactions using a flaviviral system theorized to contain crucial intragenomic RNA-RNA interactions, Japanese encephalitis virus (JEV).

JEV is a mosquito-borne flavivirus in the genus Flavivirus (family Flaviviridae), which contains several pathogenic viruses such as Dengue virus (DENV), West Nile virus (WNV), Zika virus (ZIKV), yellow fever virus (YFV). JEV is the leading cause of viral encephalitis in Southeast Asia and the Western Pacific, with approximately 68,000 cases globally each year (6). Currently, no approved treatments are available following infection by JEV or other flaviviral infections (7), which lends to the importance of further investigation. JEV is transmitted through competent mosquito vectors of the genera Aedes and Culex (8), suggesting that flaviviral infections will likely become more prevalent as global temperatures rise and vector populations expand (9). Like other flaviviruses, JEV is an enveloped virus with a single-stranded (+)-sense RNA genome of approximately 11,000nt in length (10). A single open reading frame encodes for a polyprotein and is flanked by highly structured 5’ and 3’ untranslated regions (UTRs) (11). The genome has a type I cap at the 5’ end (m7GpppAMP) and lacks polyadenylation at the 3’-terminus. The single open reading frame is cleaved post-translationally into three structural and seven non-structural proteins (12,13). During replication, the 5’ and 3’ terminal regions (TRs) in flaviviruses undergo long-range intragenomic RNA-RNA interactions, thereby forming a so-called panhandle structure that mediates recruitment of the viral RNA-dependent RNA polymerase (NS5) (14). Removal of the TRs has shown inhibition of viral replication (14–17). A cyclization sequence of 11nt is complementary in the 5’ and 3’ TRs, which facilitates this interaction (18). Furthermore, these terminal regions show binding with a variety of human host proteins (19), including but not limited to numerous DEAD-box helicases (20). In WNV and DENV, the 5’-3’ long-range interaction has been previously demonstrated (16–18,21,22). Genome cyclization in JEV has been computationally predicted previously (23); however, detailed experimental verification is still missing.

In this study, we explore the novel application of complementary computational and biophysical techniques that have not been used to characterize RNA-RNA interactions. Through a computational approach, we first identified an isolate of JEV that we hypothesized to interact with high affinity, then evaluated a consensus duplex structure of the 5’ and 3’ TRs of 20 different flaviviruses and measured their conservation. With this knowledge and utilizing various biophysical characterization techniques, we directly demonstrate for the first time that JEV 5’-3’ TRs interact *in vitro* with nanomolar affinity and with 1:1 stoichiometry. Furthermore, we isolated and identified the RNA-RNA complex using size exclusion chromatography coupled multi-angle light scattering (SEC-MALS). We additionally use analytical ultra-centrifugation (AUC) as an orthogonal biophysical validation as evidence of JEV 5’-3’TR interaction. We provide computational evidence which complements our experimental data showing that the cyclization sequence interaction is kinetically favorable and will out-compete potential homo-dimer RNA interactions. Finally, we present a low-resolution *ab inito* model showing the potential architectural arrangement of this RNA-RNA interaction in solution. This characterization can be used as a foundation in potential pharmaceutical therapies to inhibit viral replication through cyclization interruption and to help understand the replication pathway/mechanism by which NS5 replicates the viral genome.

## Materials and Methods

### Computational assessment of RNA-RNA interaction genome cyclization

Putative interaction sites between 3’ and 5’TRs of 109 JEV isolates were predicted with IntaRNA v3.2.0, RNAup v2.4.18, and RNAcofold (24–28). A consensus secondary structure of the 5’/ 3’ terminal regions in 20 phylogenetically related flaviviruses was computed with RNAalifold v2.4.18 (29) from the ViennaRNA Package v2.4.18 (30), based on a structural nucleotide multiple sequence alignment computed with LocARNA v2.0.0RC8 (31).

We evaluated whether the predicted TR interactions are kinetically feasible by applying a novel direct path model. In this model, the full target interaction is formed starting from an initial seed interaction, which folds into the target interaction by adding or removing target base pairs. All substructures on a folding path consist of consecutive base pairs of the target interaction. Consequently, paths are ‘direct’ since they cannot contain detours via non-target interaction base pairs. We compute the free energy of each substructure analogous to the RNAup energy model. The two main contributions are the cost of making the interaction site unpaired in the two intramolecular structures and the stabilizing contribution of the interaction base pairs. Both contributions were obtained from the ViennaRNA library. We can model interaction formation as a Markov process based on this direct paths model and the energy model/function. The overall rate at which an interaction is formed is determined by the energy barrier along the direct folding path (activation energy). Since the model’s possible substructures (aka states) can be completely described by their first base pair *i* and last base pair *j,* they form a 2D energy landscape, and any direct path from a first base pair to the full interaction can be drawn, as shown in figure S2. Plots of these energy landscapes allow for the visual identification of barriers along the folding path.

### Preparation and Purification of non-coding RNA

JEV 3’TR, 3’ TR Mut, and 5’TR constructs of JEV were designed based on the GenBank sequence of KR265316.1, while JEV 5’ and 3’ UTR were based on KT957419.1. cDNA sequences were prepared in pUC57 plasmids under the control of T7 RNA polymerase. The cDNA was flanked by two additional G nucleotides on the 5’ end and an XbaI restriction enzyme cut site (T^CTAGA) on the 3’ end. The 3’TR and 5’TR constructs are graphically represented in figure 1A to enhance clarity on which portion of the terminal region is involved in each experiment. All RNA construct sequences are listed below with an underlined region representing the theoretical cyclization sequence (CS) which forms the 11 nucleotide base pairing complement between the 5’ and 3’TR.

**Figure 1.**
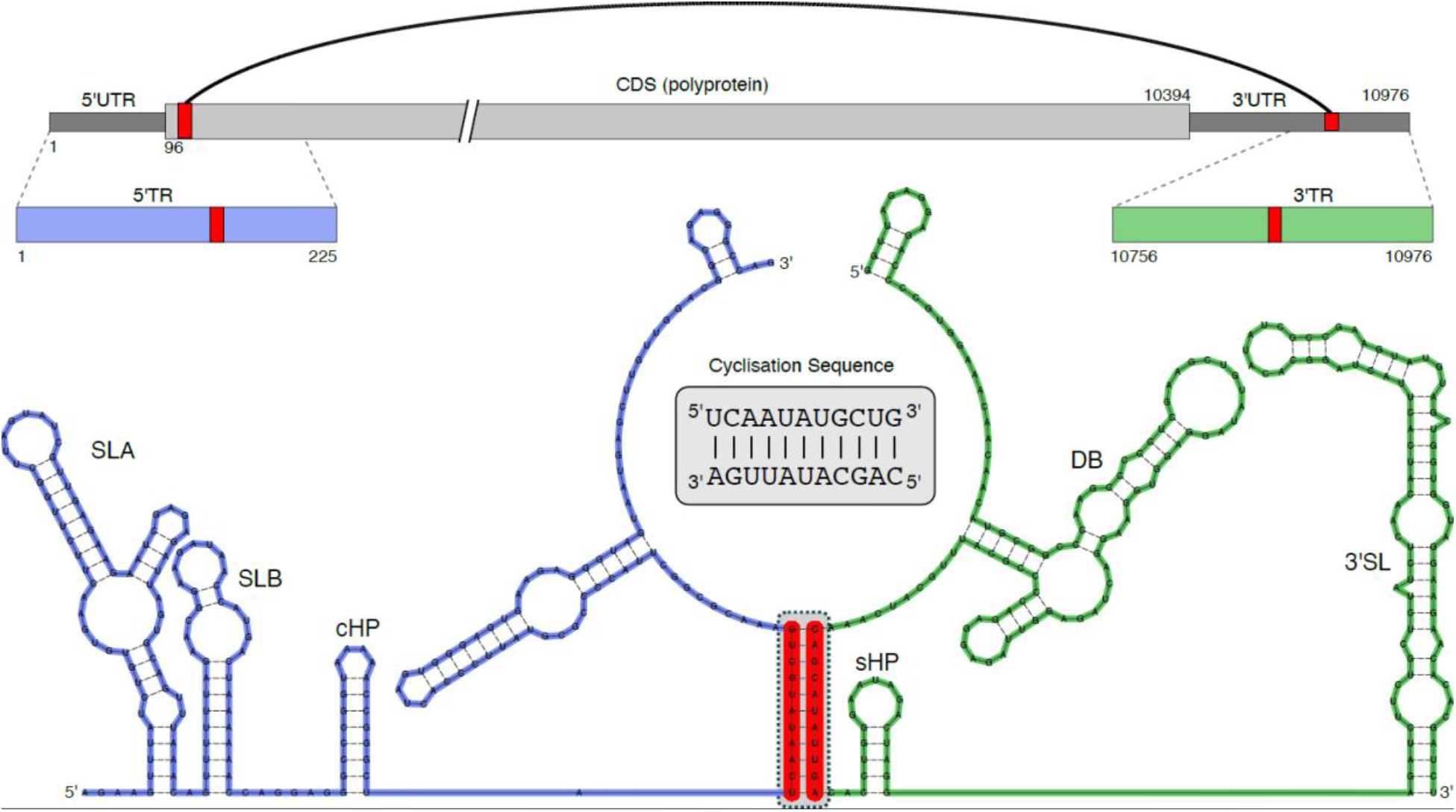
JEV structural organization and theoretical secondary structure prediction. Visual representation of JEV genome organization and different construct regions which were designed and studied. Predicted secondary structure of the cyclisation between JEV 5’TR and JEV 3’TR from constraining co-folding, highlighted red portions represents the 11 conserved nucleotides which form a stable base pairing interaction to facilitate the long-range genome interaction.

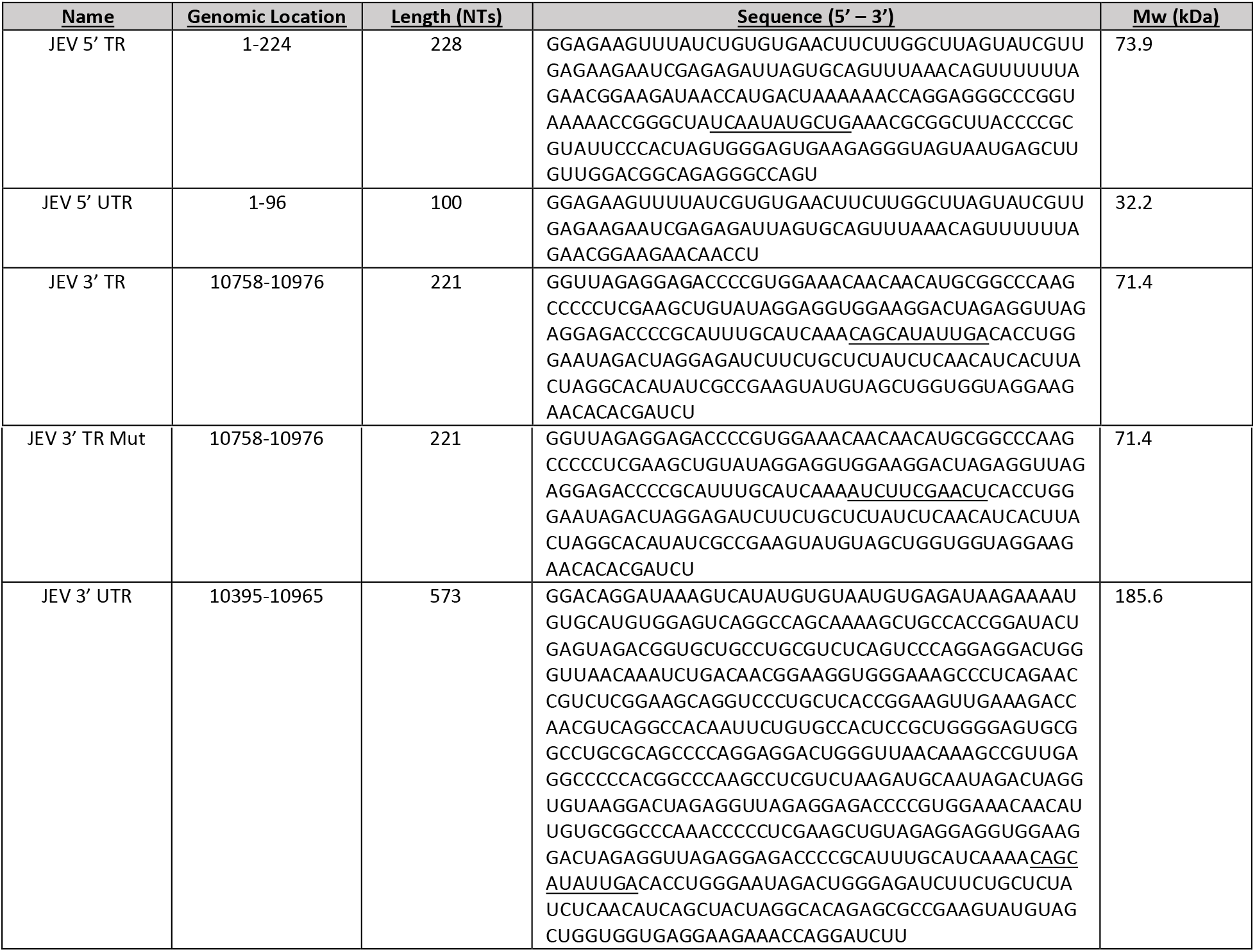

RNA was prepared via *in vitro* transcription reaction using T7 RNA polymerase (purified in-house) followed by size-exclusion chromatography (SEC) purification using a Superdex 200 Increase GL 10/300 (Global Life Science Solutions USA LLC, Marlborough, MA, USA) in JEV RNA Buffer (10 mM Bis-tris pH 5.0, 100 mM NaCl, 15 mM KCl 15 mM MgCl2, 10% glycerol) via an ÄKTA pure FPLC (Global Life Science Solutions USA LLC, Marlborough, MA, USA) with a flow rate of 0.5 mL/min. Urea-polyacrylamide gel electrophoresis (Urea-PAGE) was utilized to analyze SEC peak fractions. We mixed 10 μL of each fraction with 2 μL of denaturing RNA loading dye and loaded it into a 1.0 cm well PAGE casting plate (Bio-Rad Laboratories, Mississauga, ON, Canada). Urea-PAGE (7.5%) was run at room temperature, 300V, for 25 min (20 min for JEV 5’ TR) in 0.5x TBE (Tris-Borate-EDTA) buffer (heated), followed by staining with Sybr safe (Thermofisher Scientific, Saint-Laurant, QC, Canada) and visualization. Fractions containing a single band were deemed acceptable and used in subsequent experiments.

### Light Scattering

Multi-angle light scattering (MALS) experiments were performed on a Dawn® (Wyatt Technology Corporation, Santa Barbara, CA, USA) multi-angle light scattering instrument with 18 detector angles utilizing a 658 nm laser. Furthermore, an Optilab® (Wyatt Technology Corporation, Santa Barbara, CA, USA) refractometer was also positioned downstream to measure the solvent refractive index and absolute concentration of solutes. These instruments were positioned in line with an SEC column (Superdex 200 increase 10/300 GL, Global Life Science Solutions, USA LLC, Marlborough, MA, USA) attached to an ÄKTA pure FPLC (SEC-MALS). All experiments were performed at ambient room temperature (20°C) with the same flow rate and buffer as previous SEC experiments described above. The refractive index of the solvent was defined as 1.3308 (measured by in-line Optilab® refractometer), while the dn/dc (refractive index increment) value of 0.1720 mL/g was used for all RNAs (32). The final concentration of 3’ TR and 5’ TR RNA used was 1.3 μM and 3.5 μM, respectively, in a combined volume of 500 μL. Samples were incubated for 3 hours prior to loading. Data were analyzed using Astra v8.0.0.25, and absolute molecular weight (*M_w_*) was calculated using Equation 1 for each elution point, where: *R*(*θ*) is Rayleigh’s ratio, *K* is the polymer constant, and *c* is the concentration of the solution.

*Equation 1*

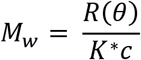

### Analytical Ultra-Centrifugation

AUC data for 5’ TR and 3’ TR were collected using a Beckman Optima AUC centrifuge and an AN50-Ti rotor at 20°C at the Canadian Centre for Hydrodynamics at the University of Lethbridge. We measured the 5’TR (214 nM, 0.5 OD @260 nm), 3’TR (224 nM, 0.5 OD @260 nm), and a 1:1 mixture of 5’TR and 3’TR samples into standard Beckman Coulter cell housings equipped with Epon-2 channel centerpieces, and fitted with sapphire windows, in JEV RNA Buffer. As a first step, we centrifuged samples at 25,000 rpm and collected scans at 20-second intervals. We used the UltraScan-III package (33) to analyze all data via supercomputer calculations in-house. We analyzed the SV-AUC data using two-dimensional spectrum analysis (2DSA) with simultaneous removal of time-invariant noise, meniscus, and bottom positions fitted, followed by enhanced van Holde-Weischet analysis (34). We estimated the buffer density and viscosity corrections with UltraScan (1.0269g/cm3 and 1.293 cP, respectively). All hydrodynamic parameters were corrected to standard conditions at 20°C and water

### Fluorescent labeling of RNA

Purified RNA was subjected to labeling at the 5’ end by the fluorophore Alexa 488 (Thermofisher Scientific, Saint-Laurant, QC, Canada). One milligram of A488 was resuspended in 175μL of 0.2M KCl. 7.5μL concentrated RNA (>100μM) was added to 1.25mg 1-ethyl-3-(3-dimethylamino) propyl carbodiimide hydrochloride (EDC) prior to the addition of 10μL resuspended A488. Samples were vortexed until contents were dissolved entirely before adding 20μL 0.1M imidazole, pH 6. Reactions were incubated in a 37°C water bath overnight in the absence of light, followed by the removal of free dye through 10 kDa Vivaspin® 500 centrifugal concentrators (Sartorius Stedim Biotech, Göttingen, Lower Saxony, Germany). After removing all free dye, labeled RNA was diluted in JEV RNA Buffer, and fluorescence checks were conducted using microscale thermophoresis (MST).

### Microscale thermophoresis

A 2-fold serial dilution was performed on the RNA ligand, either 3’TR, 3’TR Mut, or 3’UTR, where the highest concentration in the assay was 21 μM, 9.8 μM, and 5.5 μM, respectively. A constant amount of fluorescently labeled RNA Target, 5’TR, or 5’UTR, was added to each serial dilution of RNA ligand resulting in a final concentration of 25 nM and 52 nM, respectively. Mixtures were incubated at room temperature for 3 hours and then loaded into a Nanotemper Technologies Monolith® NT.115 instrument (Munich, Germany) using standard capillaries. Thermophoresis was measured at room temperature (22°C) and performed using 100% excitation power along with medium IR-Laser power. Initial fluorescence migration was measured from (−1.0 to 0 s) and used to normalize the measured fluorescent migration time (9.0 to 10.0 s). Three independent replicates were analyzed using MO.Affinity Analysis software v2.1.3 and fit to the standard K_D_ fit model, which describes a 1:1 stoichiometric molecular interaction according to the law of mass action. The dissociation constant (*K_D_*) is estimated by fitting equation 1., where *F*(*c*) is the fraction bound at a given ligand concentration c. Unbound is the *F_norm_* signal of the isolated target; Bound is the *F_norm_* signal of the complex, while *c_target_* is the final concentration of the target in the specific assay.

*Equation 2*

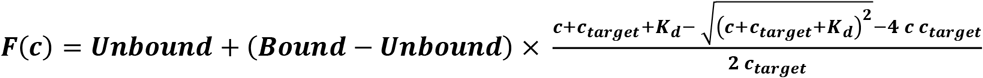

### Small-angle X-ray scattering

RNA sample data collection was performed on the B21 HPLC-SAXS beamline at Diamond Light Source (Didcot, Oxfordshire, UK), as reported elsewhere (35). An Agilent 1200 (Agilent Technologies, Stockport, UK) HPLC was utilized through connection to a specialized flow-cell, whereas 50 μL of each purified RNA (5’ TR, 3’ UTR, and 5’ TR+3’UTR respectively) were injected into a JEV RNA buffer equilibrated Shodex 403KW-4F HPLC column ( Showa Denko America Inc., New York, NY, USA) with a flow rate of 0.160 mL/min. Concentrations were 1.2 mg/mL and 1.1 mg/mL for 5’ TR and 3’ UTR, respectively, and the complex was a 1:1 volume mixture of both. Frames were exposed to synchrotron radiation (x-rays) for 3 seconds for a total of ~600 frames. The resulting data was buffer subtracted using Chromixs (36) for each sample peak, and then data analysis was performed using the ATSAS suite of programs (37). The radius of gyration (Rg) was evaluated through Guinier analysis, while additionally determining sample quality (38) and relative foldedness of RNA molecules were determined via dimensionless Kratky analysis (39). GNOM was used to perform paired distance distribution P(r) analysis to obtain real space Rg and maximum particle dimension (D_max_) measurements (40,41). Using P(r) derived information, 100 models were generated for 5’ TR and 3’ UTR using DAMMIN (42). Following simulated annealing via DAMMIN, 5’ TR models were averaged and filtered to produce a single representative model using DAMAVER and DAMFILT (42,43). Given the size of 3’ UTR, we decided to cluster representative models instead of generating a singular averaged model via DAMCLUST (43). Model reconstruction for the 5’ TR and 3’ UTR complex was generated through MONSA (42) using data input from 5’TR, 3’UTR, and data from 5’TR+3’UTR. 100 models were generated and then partitioned into representative clusters via DAMCLUST.

## Results and Discussion

### Computational analysis of the cyclization RNA-RNA interacting element in JEV and related flaviviruses

We performed computational analysis of the potential long-range interaction between the 5’ and 3’ TRs in more than 100 JEV isolates. This work revealed that all isolates interact via the canonical 11nt cyclization sequences in their 5’ and 3’ TRs. Based on this work, we selected JEV isolate KR265316.1 as a model to assess long-range RNA-RNA interactions *in silico* and *in vitro.* The Flaviviral TRs are known to harbor functional RNA elements such as stem-loops A and B in the 5’UTR, a short conserved hairpin (cHP) at the beginning of the coding region, and multiple cis-regulatory elements in the 3’UTR, such as dumbbell and terminal 3’ stem-loop (3’SL) structures (44,45). These conserved elements exert crucial roles in the flaviviral life cycle; therefore, we required them to be formed by constraint co-folding of the 5’ and 3 ‘TRs. As presented in Figure 1, the resulting duplex structure suggests that the canonical 11nt cyclization sequence in the 5’ and 3’ TRs is responsible for mediating the interaction. Furthermore, the other canonical secondary structures, such as SLA, SLB, and cHP in the 5’ TR with sHP, the 3’SL, and the 3’DB in the 3’ TR, are also present (Figure 1).

We performed a comparative genomics assay in phylogenetically related viruses to further assess flaviviral long-range RNA-RNA interactions. To this end, we analyzed the propensity of duplex formation between 5’ and 3’TRs in 20 mosquito-borne flaviviruses (Figure S2) utilizing consensus structure evaluation of the terminal genomic regions (Figure 2). Interestingly, the tendency to form a long-range interaction via the canonical cyclization motif is more pronounced in the consensus duplex than in the single sequence (JEV only) analysis. Specifically, the 11bp duplex is formed in the consensus structure without any constraints on the known 5’ and 3’ UTR elements (Figure 2). This tendency and conservation for long-range interaction in 20 sampled mosquito-borne flaviviruses reveal the importance of further understanding the RNA-RNA interaction, which mediates this cyclization interaction.

**Figure 2.**
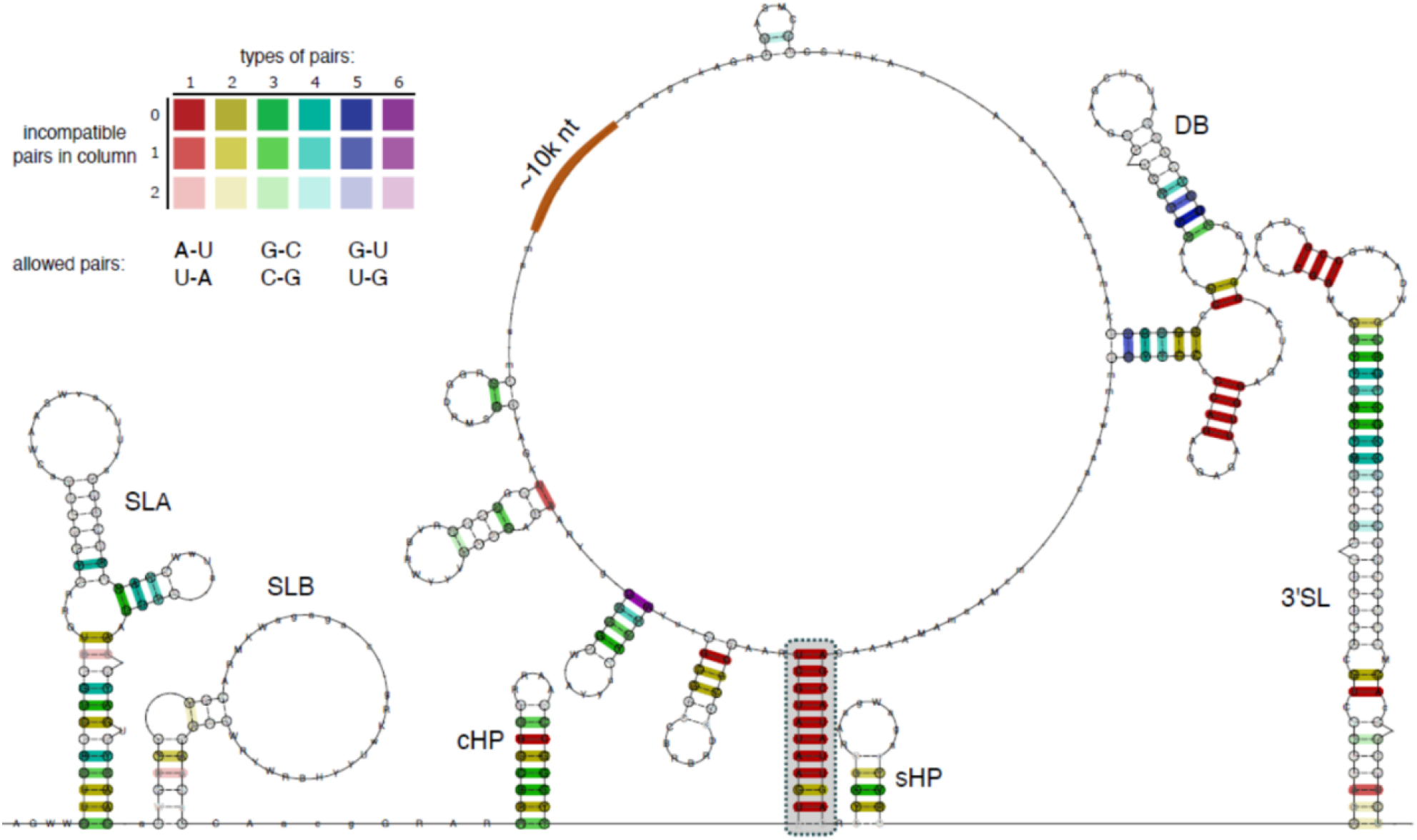
Consensus secondary structure of the 5’TR/3’TR long-range interaction, computed from a structural multiple sequence alignment of 20 mosquito-borne flaviviruses. (Figure S1). Coloring of base pairs follows the RNAalifold schema and indicates different covariation levels, ranging from red (no covariation, full primary sequence conservation) to violet (full covariation, all six possible combinations of base pairs) at corresponding columns of the underlying alignment. The duplex formed by the almost fully sequence-conserved 11 nt cyclization sequences (highlighted in gray) represents the only long-range interaction in the consensus structure. Canonical 5’TR elements (SLA, SLB, and cHP), as well as 3’TR elements (DB, sHP and 3’SL) are predicted to fold in the consensus structure, indicating that they are energetically more favourable than an extended long-range interaction duplex structure.

Additionally, a kinetic analysis of the canonical cyclization structure suggests that it is also kinetically favoured in all investigated JEV isolates. Exemplarily, the energy landscape of the cyclization structure from isolate KR265316.1 is shown in Figure 6 (discussed later). Independent of the selected start base pair, every interaction extension step leads to a more stable structure. Thus, there are no barriers along the folding paths, and the interaction can form fast. We then sought to validate the *in silico-derived* cyclization interaction in JEV through extensive biophysical characterization *in vitro.*

### In vitro transcription and purification of JEV non-coding RNA for interaction studies

JEV viral non-coding RNAs were purified immediately after *in vitro* transcription using SEC, like in previous works (20,46,47). The elution profile for 5’TR (Figure 3A) indicates that the RNA elutes at approximately 11.5 mL as a single monodispersed species, evident by the typical Gaussian distribution. 3’TR elutes as a multimodal distribution with an observable peak at ~12 mL, consistent with its size compared to 5’TR (Figure 1A). The 5’TR and 3’TR transcripts have very similar molecular weights, 73.5 kDa and 71.3 kDa, respectively, which is reflected in the chromatogram as both monomeric peaks appear to have a similar elution volume ~11.5-12.0 mL (Figure 3A). Higher-order oligomeric species for 3’TR can be observed at ~8 −11 mL, and these elution fractions we avoided for downstream experiments. To determine if the SEC elution fractions contained the correct-sized RNA species, we utilized urea-PAGE. Figure 3B (left side) shows 3’TR fractions from the right side of the ~12 mL peak containing the appropriate length RNA species (227 nt). Only selected fractions (12.5 – 13.5 mL) were pooled to avoid contamination from any oligomeric species (Figure 3A, red). 5’TR elution fractions (Figure 3B, right side) demonstrate, as expected, a single species of the correct RNA size (221 nt) across the peak. These fractions were pooled similarly to 3’TR and highlighted for clarification (Figure 3A, green). 3’TR Mut, 3’UTR, and 5’UTR (additional RNA used later) were transcribed and purified identically to the above RNA, and only fractions containing a single band were pooled and used in downstream experiments.

**Figure 3.**
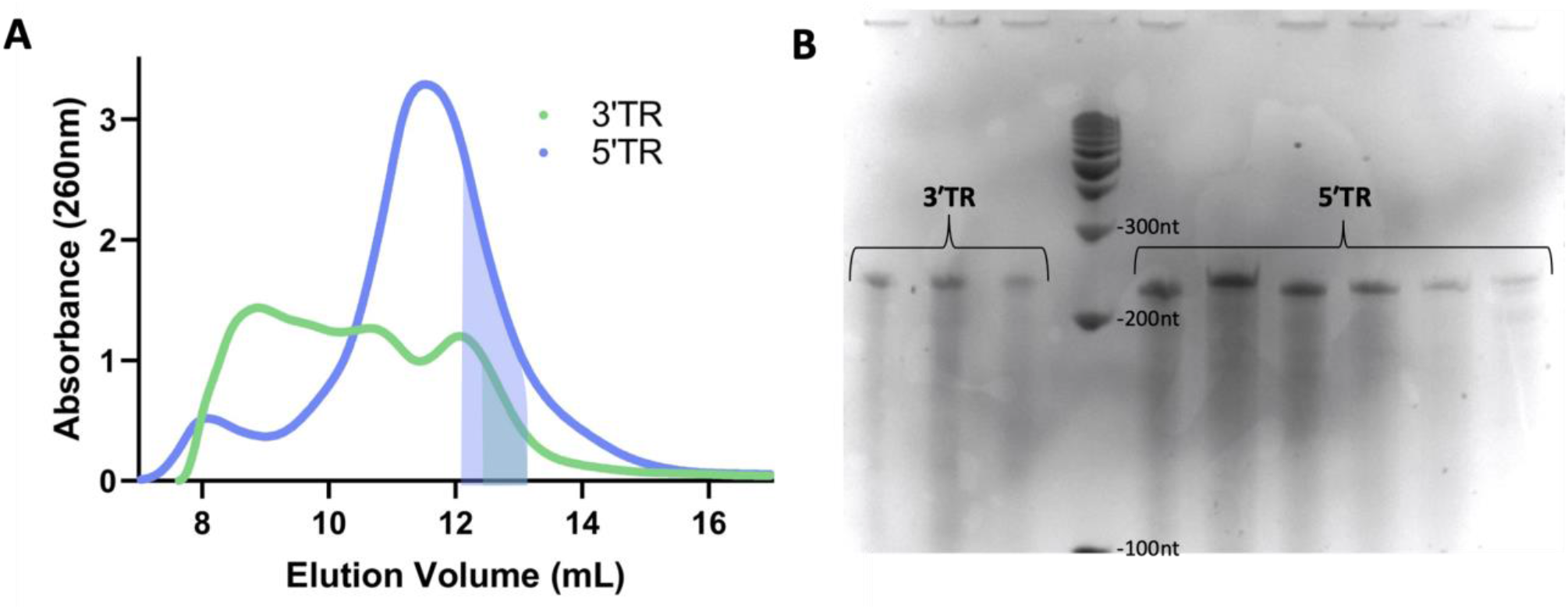
Purification of JEV TR RNA A) Size Exclusion Chromatogram representing the purification of both RNA. Shaded boxes represent the region which was collected for downstream experiments to avoid any potential oligomeric species. B) Urea PAGE of associated size exclusion chromatography fractions showing a single size of RNA (~220nt) which is the correct size of the expected RNA.

### Biophysical analysis of the RNA-RNA interacting Complex

Due to the highly conserved nature of Flavivirus TRs, it is theorized that JEV will utilize similar 3’ – 5’ long-range TR interactions as a necessary step in viral RNA amplification, like other members (49). This interaction has been shown in various Flaviviridae family members (16–18,21,22). SEC can separate biomolecules based on their sizes, and MALS allows for the absolute molecular weight determination of biomacromolecules in solution (48); we utilized SEC-MALS to investigate if the 5’ and 3’TRs form a complex. Figure 4A represents the elution profiles of JEV 3’TR, which indicates that it is almost entirely monodispersed eluting at ~12 mL with a minor peak of potential oligomeric assembly at ~11.5 mL. The elution profile of 3’ TR is consistent with our initial purification using SEC, where JEV 3’TR oligomerizes into higher-order species. JEV 5’TR also elutes at ~12 mL, consistent with previous purifications; however, it also contains a shoulder at ~11.5 mL (Figure 4A).

**Figure 4.**
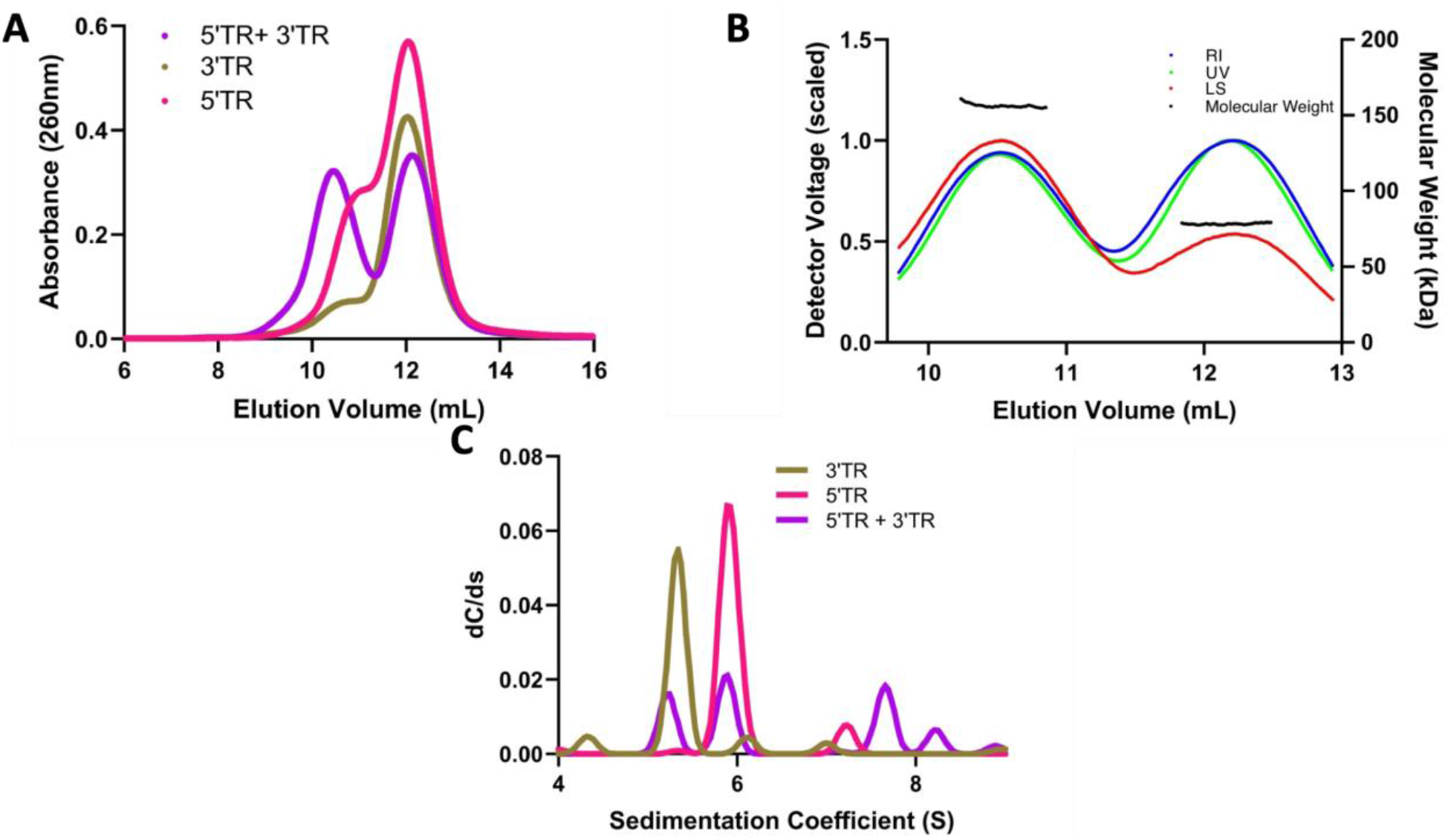
Light scattering analysis of JEV TR RNA cyclization A) Multiple size exclusion chromatography runs associated with SEC-MALS. B) MALS traces of each peak from the 5’TR +3’TR run, and the absolute molecular weight across them. C) Sedimentation distribution profiles of JEV 5’ TR and 3’TR obtained from sedimentation velocity-analytical ultracentrifugation. Sedimentation coefficient values are corrected to standard solvent conditions (20°C, water)

We then mixed both 3’TR and 5’TR and performed SEC-MALS experiments. The RNA-RNA mix resulted in a bimodal SEC chromatogram distribution, with an additional peak eluting at ~10.5 mL (Figure 4A, purple). Furthermore, we see a peak at ~12 mL where excess 5’TR RNA elutes as a monomer, consistent with the concentrations we mixed (3.5 μM of JEV 5’TR vs. 1.3 μM of JEV 3’TR). These results indicate that the TR complex has a different elution profile upon binding than any oligomeric assembly or monomeric species. Successful separation of the TR RNA complex allowed us to calculate the absolute molecular weight of each species. As presented in Figure 4B, the SEC-MALS-derived molecular weights are consistent across both peaks, indicating that both are homogenous species. Excess unbound monomer(s) elute as a single peak and are characterized at ~75 kDa, which is expected considering the similarity in their theoretical weights (73.9 kDa & 71.4 kDa). The molecular weight of the complex (~150kDa) is double that of each TR in isolation (~75kDa) and very similar to the predicted molecular weight of the complex (145.3 kDa). This change in molecular weight is indicative of a 1:1 stoichiometric interaction. This stoichiometric determination is vital for biological relevance, considering these terminal regions exist on the same genome and come together to cyclize.

AUC is a powerful biophysical technique often used to study the purity of biomolecules in solution (49). AUC subjects biomolecules to extremely high centrifugal force (up to 250,000 x g), separating them based on size, anisotropy, and density while monitoring sedimentation via an optical system. While SEC-MALS can provide us with absolute molecular weight determination, since both 5’TR and 3’TR are of similar molecular weight, we needed further validation to rule out potential RNA self-oligomerization as an explanation of the 5’TR-3’TR peak in figure 3A. AUC has been proven to be a useful technique to characterize RNA (46,47,50), but never an RNA-RNA interaction. Therefore, we utilized the specialized capabilities of AUC to provide us with an orthogonal technique that could validate our SEC-MALS results. While both 5’TR and 3’TR are similar in length, the sequences differ significantly, which will cause a change in secondary structure and, ultimately, tertiary structure. If these potential tertiary structural differences are significant enough, there should be a difference between the 5’TR and 3’TR sedimentation. Both 5’TR and 3’TR show highly monodispersed sedimentation profiles, evident by a single Gaussian distribution. 5’TR is represented by a primary sedimentation profile at ~5.9 S, while 3’TR is represented at ~5.3 S (Figure 4C). These distributions confirm that not only are the TRs homogenous, but they also sediment differently based on tertiary structure differences. This difference in sedimentation profiles allowed us to validate our previous SEC-MALS results by mixing 5’TR and 3’TR in a 1:1 ratio and performing another sedimentation velocity experiment. As with previous SEC-MALS experiments, the AUC results show an additional sedimentation peak at ~7.6 S, which is not present in the individual TR experiments allowing us to confirm JEV 5’-3’ TR interaction (Figure 4C). Additionally, since the concentrations of both RNA were ~200 nM, we can also conclude that the binding affinity (K_D_) is likely in the low nanomolar range. We believe this experiment provides the first evidence of an RNA-RNA interaction via AUC, showing that AUC can be an essential tool for showing RNA-RNA interaction while simultaneously determining sample quality and oligomeric state. Taking the AUC and SEC-MALS results together, we can confidently conclude that the additional peak(s) formed in both experiments is the complex of 5’TR and 3’TR, which provides experimental validation of computational predictions.

### Determining the RNA-RNA interaction affinity

With the above-mentioned relative binding affinity and direct evidence of interaction, we sought to quantify the affinity of the 5’ and 3’ TRs using MST. MST allows for studying the interaction between a serially diluted ligand and a fluorescently labeled target by measuring the change in fluorescent migration following excitation by an infrared laser (51). Using the difference between the “cold” and “hot” areas of the MST traces (Figure 5A), MST can determine the K_D_ of the biomolecular interaction. Additionally, figure 5A demonstrates that aggregation of the target was not observed (52) and that monomeric fractions were utilized in the assay. Our analysis indicates that the TRs interact with a K_D_ of 60 ± 9 nM (Figure 5B), which agrees with the previously reported K_D_ of 32 ± 1 nM found in WNV using isothermal titration calorimetry (22), which is reasonably similar, validating our findings.

**Figure 5.**
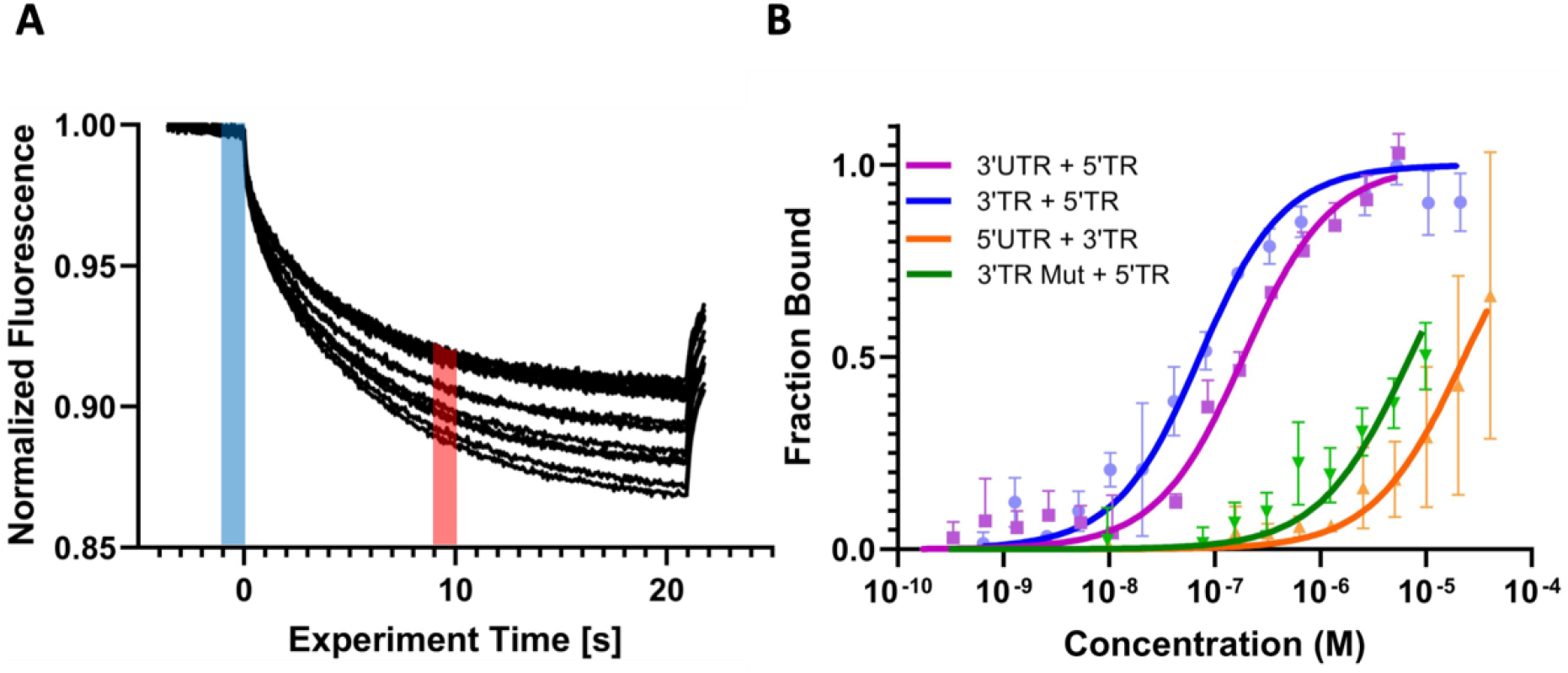
Affinity analysis of JEV TR RNA cyclization A) MST raw data traces for JEV 3’TR + 5’TR. Blue line represents the “cold” time and the red line represent the “hot” region, and the difference between the two is used to calculate the ΔF_norm_ B) Microscale thermophoresis measurements of different combinations of JEV TRs representing concentration vs fraction bound. Measurement ran on “high” MST power.

**Figure 6.**
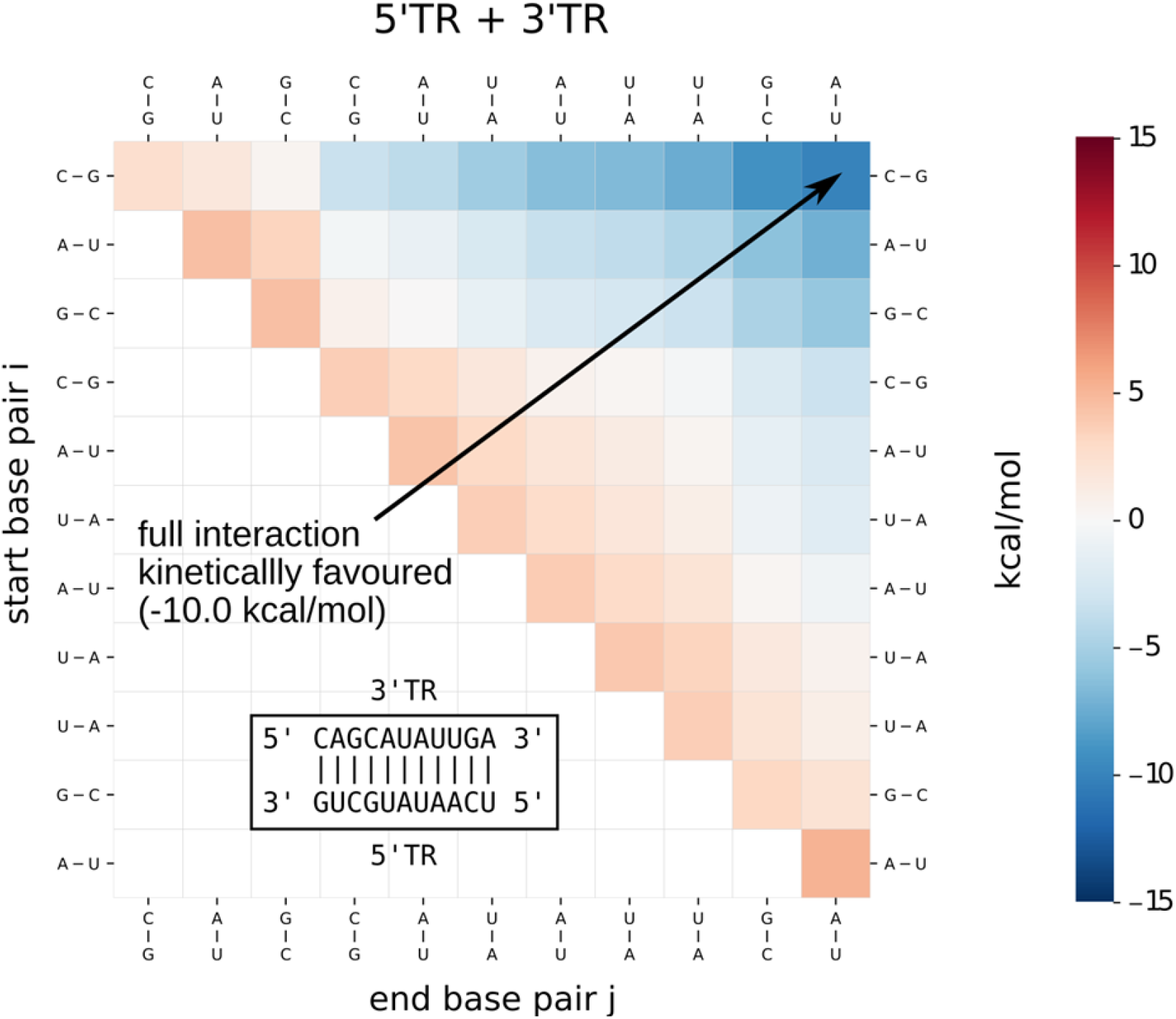
Energy landscape of the predicted 3’TR and 5’TR CS interaction. The energy landscape represents all possible substructures that can occur along a direct path from a single base pair interaction to the full CS interaction.

Consequently, the binding affinity further validates our AUC experiment, performed with ~200 nM concentrations of JEV TR RNA. To our knowledge, this is the first evidence of an RNA-RNA interaction characterized using MST, highlighting the versatility of the technique. Next, we needed to validate whether the CS is the primary driver of this RNA-RNA interaction. We computationally predicted the RNA duplex secondary structure (Figure S3), showing numerous potential base-pairing sites across the interaction, but the longest uninterrupted stretch of nucleotides was the CS. We investigated the degree to which the inclusion of the CS impacts binding affinity by exploring various fragments of the JEV TRs. JEV 3’UTR, the full-length 3’ non-coding RNA at 573 nt, and JEV 5’TR, which both still include the CS, were assayed and found to interact with a K_D_ of 169 ± 18 nM (Figure 5B). As expected, the theoretically determined highest binding isolates of JEV (5’TR and 3’TR) bound with a slightly higher affinity than the 3’UTR and 5’TR but are within the same magnitude. When excluding the CS through a truncation (JEV 5’UTR-97 nt), the interaction was almost non-existent with a K_D_ of 23 ± 6 μM. Finally, we performed the same assay with a mutant version of 3’TR, 3’TR Mut. This construct still contains both the CS and DAR sequence elements, while JEV 5’ UTR is truncated, missing both the CS and the DAR element. Therefore, we believed that the inclusion of the additional long-range binding element would result in a binding tighter than the 5’ UTR-3’TR interaction, but still considerably weaker than the canonical CS included. The interaction was as expected and was significantly lower than the 5’TR-3’TR interaction, with a K_D_ of 7 ± 1.4 μM (figure 5B). This change in affinity demonstrates that while the theoretical duplex interaction may have considerable base pairing, the primary driver of the interaction is the cyclization sequence.

### RNA-RNA Kinetic and thermodynamic Studies

As observed in Figure 3A and Figure 4A, both TRs are likely to self-associate even though careful consideration was taken to avoid this. Therefore, we utilized computational analysis to explain the self-association of TRs potentially. While our predictions find potential homodimer interactions with similar thermodynamic stability as the hetero duplex (canonical CS), these homodimer interactions are quite extended and would require extensive refolding. Based on a kinetic analysis (Figure S2), we assume that homodimers would form only short, less stable interactions, which would readily dissociate in favour of the more stable heterodimer (3’ – 5’ complex) when mixed.

To help explain our MST observations, we compared the measured dissociation constants to the predicted interactions for the different TR and UTR constructs. We calculated binding free energies at 37 °C either from the measured dissociation constants or from predictions using RNAup (predictions are in parentheses). For the three constructs, the 5’TR and 3’TR, 5’TR and 3’UTR, and 5’UTR and 3’TR complex, we obtained ΔGs of −10.2 (−10.1) kcal/mol, −9.6 (−7.6) kcal/mol and −6.6 (−7.2) kcal/mol, respectively. The more considerable discrepancy for the second construct (5’TR and 3’UTR) can be explained because the thermodynamic model predicts a refolding (compared to the known consensus structure) when the longer 3’UTR is used, occluding the CS. We also investigated potential additional interaction sites to the known CS. The most promising is a kinetically favourable interaction corresponding to the known upstream AUG Region (UAR) (53) with a predicted stability of −3.6 kcal/mol. This interaction could stabilize the CS interaction but is not strong enough to explain the duplex formation alone.

### 3-dimensional RNA-RNA interaction analysis through small-angle X-ray scattering

Having characterized the cyclization RNA-RNA binding affinity, we turned to small-angle X-ray scattering to understand the 3-dimensional interaction in solution. Using SEC-SAXS, it is possible to differentiate species based on size and shape prior to actual SAXS measurements, providing increased confidence in monodispersity and allowing for the characterization of a potential complex from two mixed samples (54–57). MST experiments revealed that the binding affinity of the 5’TR and 3’ UTR were comparable to the 5’TR-3’TR interaction, so we decided to perform structural experiments on the larger RNA construct (3’ UTR) to gain a better characterization of how this RNA interaction is arranged in solution and its relative flexibility.

Three SAXS data sets were collected: 5’ TR, 3’ UTR, and 5’TR + 3’UTR. Each was merged and represented in figure S4a as relative intensity vs. scattering angle. Guinier analysis was performed on each data set to determine the radius of gyration (Rg) for each RNA resulting in 71.61 ± 0.106 Å and 111.5 ± 0.33 Å for 5’TR and 3’UTR respectively (fig S4b). Complex (5’TR + 3’UTR) Guinier analysis reveals an Rg of 127.6 ± 0.34 Å, an increase compared to each RNA. The linear regression of each sample demonstrates that each sample is monodispersed and free of any electrostatic interactions between similar molecules (58–60). Intensity data were transformed into dimensionless Kratky analysis data (Fig S4c) to evaluate approximate foldedness and conformation (61). Kratky analysis (fig S4c) for all three RNA suggests extended conformations of folded RNA based on the relative plateauing of each data set (58). Finally, each data set was evaluated using paired-distance distribution (p(r)) analysis in which reciprocal space data is converted into real-space electron density data via indirect Fourier transformations (62). P(r) analysis presents real space R_g_ values of 71.8 ± 0.104 Å, 112.0 ± 0.329 Å, and 128.1 ± 0.355 Å for the 5’TR, 3’UTR, and the complex, respectively (fig S4d). Real-space Rg values have a very high level of agreement with the reciprocal-space values, indicating the validity of the entire data set. Furthermore, the shape of each P(r) plot indicates an elongated conformation, whereas a gaussian distribution with a peak at D_max_/2 would be indicative of a spherical, globular molecule. The maximum distance for each RNA data set was 224 Å, 345 Å, and 400 Å for the 5’TR, 3’ UTR, and complex, respectively. Notably, the D_max_ values illustrate that while the complex is larger than either individual RNA, likely that the interaction is not end-to-end. The maximal distance for an end-to-end interaction would be considerably larger than 400 Å, suggesting there may be considerable overlap between the 5’TR and 3’UTR.

*Ab inito* modeling through DAMMIN was then performed on 5’TR and 3’UTR data sets to generate low-resolution three-dimensional structures. 100 models were generated for each RNA showing favorable agreement via χ^2^ values of 1.15 and 1.12 for the 5’TR and 3’UTR, respectively. The 5’ TR was filtered and merged into a singular representative structure with a normalized spatial discrepancy (NSD) of 0.937 ± 0.020, indicating a good fit of each structure to the representative filtered model (figure 7A)(63). Represented in blue, the 5’ TR is an elongated RNA structure like other RNA of similar size (47,50,64). The 3’UTR, however, could not be filtered and averaged together to get a singular representative model, even though the average χ^2^ value was 1.12, suggesting that each model strongly agreed with the original scattering data. We, therefore, chose to cluster the 100 respective models from the 3’UTR, resulting in 22 distinctly different clusters represented by the pie chart in figure 7B. This suggests that the 3’UTR has flexible regions, similar to other long non-coding RNA (65,66). The three largest clusters of models are color coded to the pie chart, and while each is classified as a distinct structure, they contain a similar overall architecture: each having a large center pocket and large portions of electron density towards each terminus.

**Figure 7.**
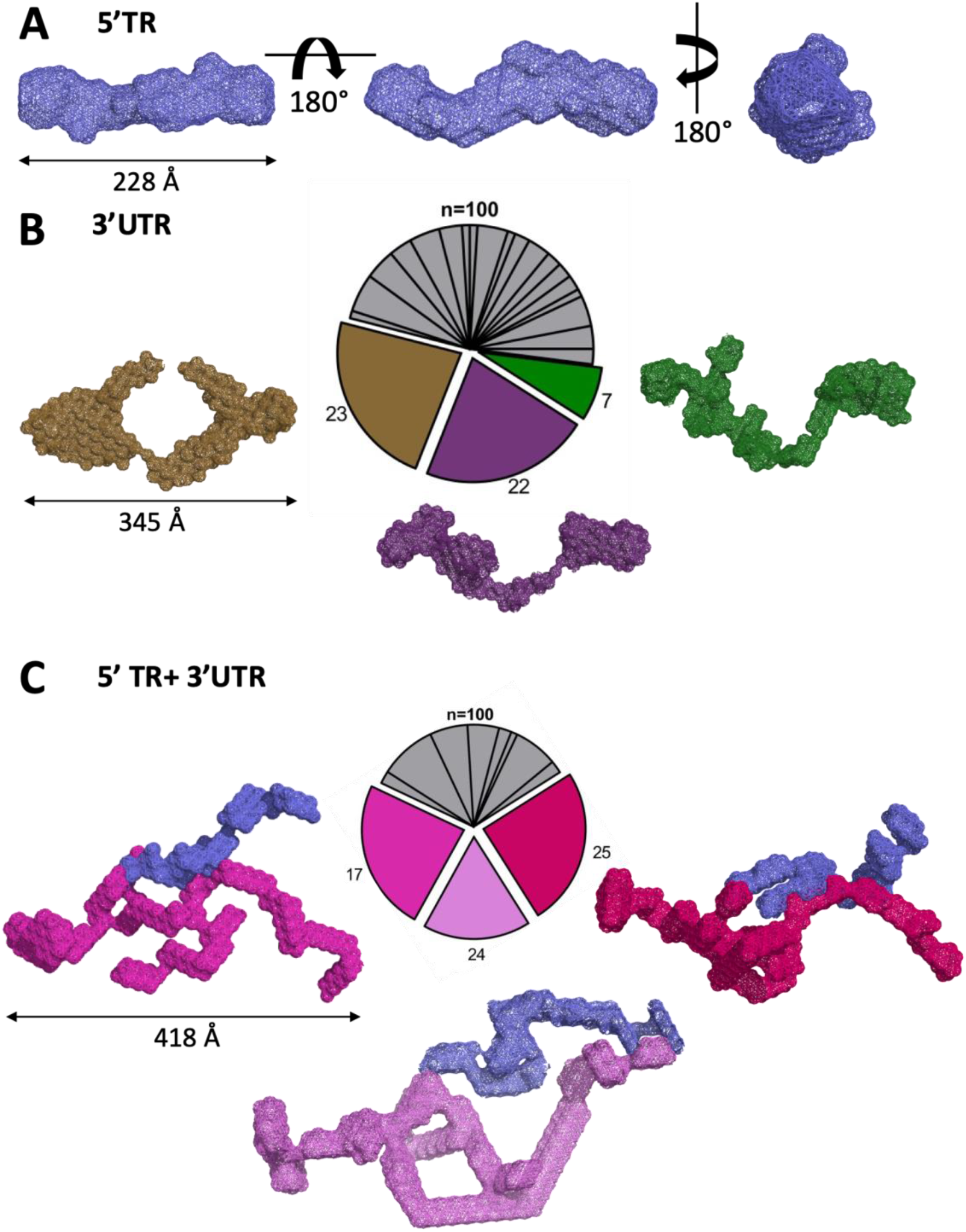
Small-angle X-ray scattering *ab inito* model reconstructions of JEV RNA. A) JEV 5’ TR RNA, rotations are 180° about both the X and Y axis and showing D_max_. B) JEV 3’ UTR RNA illustrating relative conformational fluidity through clustering analysis of 100 *ab inito* models. Shown models represent three of the largest clusters of models. C) JEV 5’TR+3’UTR RNA-RNA complex. JEV 5’TR is represented in blue, while 3’UTR is represented in shades of red/purple.

Importantly, since the affinity of the 5’TR – 3’UTR interaction was sufficiently low, we believed that the complex would remain intact throughout SEC-SAXS allowing for peak separation and, ultimately, data collection. Creation of the three-dimensional models required multiple distinct inputs, differing from the terminal regions individually. Using MONSA, multi-phase bead modeling can simultaneously fit multiple scattering curves to a single data set (42,67). Therefore, discerning which portion of the total scattering data was contributed by each RNA element required inputting data collected from the 5’TR, the 3’TR, and the complex. Importantly, when evaluating χ^2^ values for the complex, each input is scored based on its agreement with the raw scattering data with approximate values of 1.20, 1.40, and 1.32 for the 5’TR, 3’UTR, and complex, respectively. These values show that each distinct scattering curve fits into the data set and has a good agreement. Like the 3’UTR on its own, a singular averaged model cannot accurately represent the RNA, which led us to cluster the model representations (Figure 7C) similarly. This clustering resulted in 11 distinct representations of the solution structure, half as many clusters as the 3’UTR alone, with 66% of the total structures falling into 3 clusters. Clustering provides evidence that the interaction causes a change in the conformational space and a subsequent reduction in the flexibility of the 3’UTR. Figure 7C represents the top 3 conformational clusters, with the 5’TR in a consistent blue color and the 3’UTR in pink/red. The 5’TR remains relatively unchanged upon interaction with the 3’UTR, still adopting an elongated shape similar to its previous conformation (Figure 7A). Most of the conformational change to the complex is changes to the 3’UTR. Overall, the central pocket, which was visible in the top 3’UTR conformations, is no longer visible, and the electron density has been re-distributed from being concentrated around the ends of the molecule (see figure 7B). Furthermore, the 3-dimensional complex shows that there is likely not considerable 5’-3’ duplex formation, with generally 1-2 contact points, one of which is likely the CS. Structural information about this complex can also inform on the potential mechanism by which the flaviviral RNA-dependent RNA polymerase NS5 interacts with the 5’ TR SLA yet transcribes the negative sense genome 3’ to 5’ (14,68–70). However, the nature of SAXS being low-resolution means we cannot accurately pinpoint specific RNA motifs such as the CS or the 3’ or 5’ ends. Future directions will need to be focused on gaining a higher-resolution model of the interaction through either computational modeling with SAXS envelopes or cryo-EM imaging.

### Conclusion

Our study demonstrates that combining computational and biophysical approaches can provide detailed insights into RNA-RNA interactions that govern many fundamental biological processes. Our SEC-MALS, AUC, and MST data also suggest, for the first time, that these techniques can be used to study RNA-RNA interactions in solution. We demonstrate that computational analysis, such as kinetic landscapes, can provide vital evidence for predicting RNA-RNA interactions and even sites of interactions. Additionally, our binding affinity data presents that the CS is the primary driver of terminal region interaction with a nanomolar affinity. This binding affinity is critical to developing potential inhibitory therapeutics targeting the cyclization sequence. Additionally, we present the 3-dimensional low-resolution structure complex structure in solution, providing evidence that it is likely flexible and dynamic, with a small amount of duplex formation between the 5’ TR and 3’UTR. Our work directly contributes to understanding the Flaviviral cyclization conservation that could help to developing therapeutics with a potential for multi-virus inhibition.

## Data availability

JEV isolates for computational analysis:

We selected complete JEV isolates (including 3’ and 5’ UTR) from NCBI GenBank and used the first 225 nts (5’TR) and the last 221 nts (3’TR) to perform interaction predictions. The corresponding accession numbers can be found in the ‘JEV.txt ‘ supplemental file

## Funding and additional information

T.M. is supported by a Natural Sciences and Engineering Research Council (NSERC) PGS-D award. S.M.P was supported by NSERC Undergraduate Student Research Award, S.T and AD are supported by the NSERC Discovery grant awarded to T.R.P., and C.R.N. was supported by Alberta Innovates Graduate student award. M.W. is supported by the Austrian Science Fund (FWF I-2874-N28, DK RNA Biology, F 80 RNAdeco awarded to ILH). A.H is supported through by an NSERC CGS-D award. This research was funded by the NSERC Discovery grant, RGPIN-2017-04003 to T.R.P., T.R.P. is a Canada Research Chair in RNA and Protein Biophysics. Infrastructure support to T.R.P. was provided by the Canada Foundation for Innovation and NSERC RTI Grants. This work was supported by the Canada 150 Research Chairs program (C150-2017-00015, BD), the Canada Foundation for Innovation (CFI-37589, BD), the National Institutes of Health (1R01GM120600, BD) and the Canadian Natural Science and Engineering Research Council (DG-RGPIN-2019-05637, BD). UltraScan supercomputer calculations were supported through NSF/XSEDE grant TG-MCB070039N (BD). This research was partially funded by Austrian Science Fund (FWF) project I 6440-N to M.T.W.

## References

1. Zhang, F. and Lupski, J.R. (2015) Non-coding genetic variants in human disease. Hum Mol Genet, 24, R102–110.

2. Balas, M.M., Hartwick, E.W., Barrington, C., Roberts, J.T., Wu, S.K., Bettcher, R., Griffin, A.M., Kieft, J.S. and Johnson, A.M. (2021) Establishing RNA-RNA interactions remodels lncRNA structure and promotes PRC2 activity. Science Advances, 7, eabc9191.

3. Aguilar, R., Spencer, K.B., Kesner, B., Rizvi, N.F., Badmalia, M.D., Mrozowich, T., Mortison, J.D., Rivera, C., Smith, G.F., Burchard, J. et al. (2022) Targeting Xist with compounds that disrupt RNA structure and X inactivation. Nature, 604, 160–166.

4. Jain, A. and Vale, R.D. (2017) RNA phase transitions in repeat expansion disorders. Nature, 546, 243–247.

5. Van Treeck, B. and Parker, R. (2018) Emerging Roles for Intermolecular RNA-RNA Interactions in RNP Assemblies. Cell, 174, 791–802.

6. Heffelfinger, J.D., Li, X., Batmunkh, N., Grabovac, V., Diorditsa, S., Liyanage, J.B., Pattamadilok, S., Bahl, S., Vannice, K.S., Hyde, T.B. et al. (2017) Japanese Encephalitis Surveillance and Immunization - Asia and Western Pacific Regions, 2016. MMWR Morb Mortal Wkly Rep, 66, 579–583.

7. Wang, Q.-Y. and Shi, P.-Y. (2015) Flavivirus Entry Inhibitors. ACS Infectious Diseases, 1, 428–434.

8. Huang, Y.J., Higgs, S., Horne, K.M. and Vanlandingham, D.L. (2014) Flavivirus-mosquito interactions. Viruses, 6, 4703–4730.

9. Tjaden, N.B., Caminade, C., Beierkuhnlein, C. and Thomas, S.M. (2018) Mosquito-Borne Diseases: Advances in Modelling Climate-Change Impacts. Trends in Parasitology, 34, 227–245.

10. Lindenbach, B., Thiel, H.J. and Rice, C.M. (2007) Flaviviridae: The viruses and their replication. Fields Virology, 1101–1151.

11. Ng, W.C., Soto-Acosta, R., Bradrick, S.S., Garcia-Blanco, M.A. and Ooi, E.E. (2017) The 5’ and 3’ Untranslated Regions of the Flaviviral Genome. Viruses, 9, 137.

12. Bollati, M., Alvarez, K., Assenberg, R., Baronti, C., Canard, B., Cook, S., Coutard, B., Decroly, E., de Lamballerie, X., Gould, E.A. et al. (2010) Structure and functionality in flavivirus NS-proteins: perspectives for drug design. Antiviral research, 87, 125–148.

13. Mandl, C.W., Holzmann, H., Kunz, C. and Heinz, F.X. (1993) Complete Genomic Sequence of Powassan Virus: Evaluation of Genetic Elements in Tick-Borne versus Mosquito-Borne Flaviviruses. Virology, 194, 173–184.

14. Filomatori, C.V., Lodeiro, M.F., Alvarez, D.E., Samsa, M.M., Pietrasanta, L. and Gamarnik, A.V. (2006) A 5’ RNA element promotes dengue virus RNA synthesis on a circular genome. Genes Dev, 20, 2238–2249.

15. You, S. and Padmanabhan, R. (1999) A Novel in Vitro Replication System for Dengue Virus: INITIATION OF RNA SYNTHESIS AT THE 3’-END OF EXOGENOUS VIRAL RNA TEMPLATES REQUIRES 5’- AND 3’-TERMINAL COMPLEMENTARY SEQUENCE MOTIFS OF THE VIRAL RNA*. Journal of Biological Chemistry, 274, 33714–33722.

16. Zhang, B., Dong, H., Stein, D.A., Iversen, P.L. and Shi, P.-Y. (2008) West Nile virus genome cyclization and RNA replication require two pairs of long-distance RNA interactions. Virology, 373, 1–13.

17. Alvarez, D.E., Lodeiro, M.F., Ludueña, S.J., Pietrasanta, L.I. and Gamarnik, A.V. (2005) Long-range RNA–RNA interactions circularize the dengue virus genome. Journal of virology, 79, 6631–6643.

18. Hahn, C.S., Hahn, Y.S., Rice, C.M., Lee, E., Dalgarno, L., Strauss, E.G. and Strauss, J.H. (1987) Conserved elements in the 3’ untranslated region of flavivirus RNAs and potential cyclization sequences. J Mol Biol, 198, 33–41.

19. Meier-Stephenson, V., Mrozowich, T., Pham, M. and Patel, T.R. (2018) DEAD-box helicases: the Yin and Yang roles in viral infections. Biotechnology & genetic engineering reviews, 34, 3–32.

20. Nelson, C., Mrozowich, T., Gemmill, D.L., Park, S.M. and Patel, T.R. (2021) Human DDX3X Unwinds Japanese Encephalitis and Zika Viral 5’ Terminal Regions. Int J Mol Sci, 22, 413.

21. Men, R., Bray, M., Clark, D., Chanock, R.M. and Lai, C.J. (1996) Dengue type 4 virus mutants containing deletions in the 3’ non-coding region of the RNA genome: analysis of growth restriction in cell culture and altered viremia pattern and immunogenicity in rhesus monkeys. Journal of virology, 70, 3930–3937.

22. Deo, S., Patel, T.R., Chojnowski, G., Koul, A., Dzananovic, E., McEleney, K., Bujnicki, J.M. and McKenna, S.A. (2015) Characterization of the termini of the West Nile virus genome and their interactions with the small isoform of the 2’ 5’-oligoadenylate synthetase family. Journal of structural biology, 190, 236–249.

23. Thurner, C., Witwer, C., Hofacker, I.L. and Stadler, P.F. (2004) Conserved RNA secondary structures in Flaviviridae genomes. Journal of General Virology, 85, 1113–1124.

24. Mann, M., Wright, P.R. and Backofen, R. (2017) IntaRNA 2.0: enhanced and customizable prediction of RNA-RNA interactions. Nucleic acids research, 45, W435–W439.

25. Busch, A., Richter, A.S. and Backofen, R. (2008) IntaRNA: efficient prediction of bacterial sRNA targets incorporating target site accessibility and seed regions. Bioinformatics, 24, 2849–2856.

26. Mückstein, U., Tafer, H., Hackermüller, J., Bernhart, S.H., Stadler, P.F. and Hofacker, I.L. (2006) Thermodynamics of RNA–RNA binding. Bioinformatics, 22, 1177–1182.

27. Hofacker, I.L., Fontana, W., Stadler, P.F., Bonhoeffer, L.S., Tacker, M. and Schuster, P. (1994) Fast folding and comparison of RNA secondary structures. Monatshefte für Chemie / Chemical Monthly, 125, 167–188.

28. Bernhart, S.H., Tafer, H., Mückstein, U., Flamm, C., Stadler, P.F. and Hofacker, I.L. (2006) Partition function and base pairing probabilities of RNA heterodimers. Algorithms for Molecular Biology, 1, 3.

29. Bernhart, S.H., Hofacker, I.L., Will, S., Gruber, A.R. and Stadler, P.F. (2008) RNAalifold: improved consensus structure prediction for RNA alignments. BMC Bioinformatics, 9, 474.

30. Lorenz, R., Bernhart, S.H., Höner zu Siederdissen, C., Tafer, H., Flamm, C., Stadler, P.F. and Hofacker, I.L. (2011) ViennaRNA Package 2.0. Algorithms for Molecular Biology, 6, 26.

31. Will, S., Reiche, K., Hofacker, I.L., Stadler, P.F. and Backofen, R. (2007) Inferring Noncoding RNA Families and Classes by Means of Genome-Scale Structure-Based Clustering. PLOS Computational Biology, 3, e65.

32. Pam Wang, R.A., Michelle Chen, Kristine Legaspi. (2020), Wyatt Technologies, pp. 1–4.

33. Demeler, B. and Gorbet, G.E. (2016) In Uchiyama, S., Arisaka, F., Stafford, W. F. and Laue, T. (eds.), Analytical Ultracentrifugation: Instrumentation, Software, and Applications. Springer Japan, Tokyo, pp. 119–143.

34. Demeler, B. and van Holde, K.E. (2004) Sedimentation velocity analysis of highly heterogeneous systems. Analytical Biochemistry, 335, 279–288.

35. Meier, M., Moya-Torres, A., Krahn, N.J., McDougall, M.D., Orriss, G.L., McRae, E.K.S., Booy, E.P., McEleney, K., Patel, T.R., McKenna, S.A. et al. (2018) Structure and hydrodynamics of a DNA G-quadruplex with a cytosine bulge. Nucleic acids research, 46, 5319–5331.

36. Panjkovich, A. and Svergun, D.I. (2017) CHROMIXS: automatic and interactive analysis of chromatography-coupled small-angle X-ray scattering data. Bioinformatics, 34, 1944–1946.

37. Manalastas-Cantos, K., Konarev, P., Hajizadeh, N., Kikhney, A., Petoukhov, M., Molodenskiy, D., Panjkovich, A., Mertens, H., Gruzinov, A., Borges, C. et al. (2021) ATSAS 3.0: Expanded functionality and new tools for small-angle scattering data analysis. Journal of Applied Crystallography, 54, 343–355.

38. Putnam, C.D. (2016) Guinier peak analysis for visual and automated inspection of small-angle X-ray scattering data. J Appl Crystallogr, 49, 1412–1419.

39. Burke, J.E. and Butcher, S.E. (2012) Nucleic acid structure characterization by small angle X-ray scattering (SAXS). Curr Protoc Nucleic Acid Chem, Chapter 7, Unit 7.18.

40. Semenyuk, A.V. and Svergun, D.I. (1991) GNOM–a program package for small-angle scattering data processing. Journal of Applied Crystallography, 24, 537–540.

41. Svergun, D.I. (1992) Determination of the regularization parameter in indirect-transform methods using perceptual criteria. Journal of Applied Crystallography, 25, 495–503.

42. Svergun, D.I. (1999) Restoring Low Resolution Structure of Biological Macromolecules from Solution Scattering Using Simulated Annealing. Biophysical journal, 76, 2879–2886.

43. Volkov, V.V. and Svergun, D.I. (2003) Uniqueness of ab initio shape determination in small-angle scattering. Journal of Applied Crystallography, 36, 860–864.

44. Ochsenreiter, R., Hofacker, I.L. and Wolfinger, M.T. (2019) Functional RNA Structures in the 3’UTR of Tick-Borne, Insect-Specific and No-Known-Vector Flaviviruses. Viruses, 11, 298.

45. Michael T. Wolfinger, R.O., Ivo L. Hofacker. (2021) In Marz, D. F. a. M. (ed.), Virus Bioinformatics. 1 ed. Chapman and Hall/CRC, pp. 36.

46. Nelson, C.R., Mrozowich, T., Park, S.M., D’souza, S., Henrickson, A., Vigar, J.R.J., Wieden, H.-J., Owens, R.J., Demeler, B. and Patel, T.R. (2021) Human DDX17 Unwinds Rift Valley Fever Virus Non-Coding RNAs. Int J Mol Sci, 22, 54.

47. Mrozowich, T., Henrickson, A., Demeler, B. and Patel, T.R. (2020) Nanoscale Structure Determination of Murray Valley Encephalitis and Powassan Virus Non-Coding RNAs. Viruses, 12, 190.

48. Wyatt, P.J. (1993) Light scattering and the absolute characterization of macromolecules. Analytica Chimica Acta, 272, 1–40.

49. Patel, T.R., Winzor, D.J. and Scott, D.J. (2016) Analytical ultracentrifugation: A versatile tool for the characterisation of macromolecular complexes in solution. Methods, 95, 55–61.

50. D’Souza, M.H., Mrozowich, T., Badmalia, M.D., Geeraert, M., Frederickson, A., Henrickson, A., Demeler, B., Wolfinger, M.T. and Patel, T.R. (2022) Biophysical characterisation of human LincRNA-p21 sense and antisense Alu inverted repeats. Nucleic acids research, 50, 5881–5898.

51. Wienken, C.J., Baaske, P., Rothbauer, U., Braun, D. and Duhr, S. (2010) Protein-binding assays in biological liquids using microscale thermophoresis. Nature Communications, 1, 100.

52. Mrozowich, T., MeierStephenson, V. and Patel, T.R. (2019) Microscale thermophoresis: warming up to a new biomolecular interaction technique. The Biochemist, 41, 8–12.

53. Friebe, P., Shi, P.-Y. and Harris, E. (2011) The 5’ and 3’ downstream AUG region elements are required for mosquito-borne flavivirus RNA replication. Journal of virology, 85, 1900–1905.

54. Brosey, C.A. and Tainer, J.A. (2019) Evolving SAXS versatility: solution X-ray scattering for macromolecular architecture, functional landscapes, and integrative structural biology. Curr Opin Struct Biol, 58, 197–213.

55. Pérez, J. and Vachette, P. (2017) In Chaudhuri, B., Muñoz, I. G., Qian, S. and Urban, V. S. (eds.), Biological Small Angle Scattering: Techniques, Strategies and Tips. Springer Singapore, Singapore, pp. 183–199.

56. O’Brien, D.P., Brier, S., Ladant, D., Durand, D., Chenal, A. and Vachette, P. (2018) SEC-SAXS and HDX-MS: A powerful combination. The case of the calcium-binding domain of a bacterial toxin. Biotechnol Appl Biochem, 65, 62–68.

57. Graewert, M.A., Da Vela, S., Gräwert, T.W., Molodenskiy, D.S., Blanchet, C.E., Svergun, D.I. and Jeffries, C.M. (2020) Adding Size Exclusion Chromatography (SEC) and Light Scattering (LS) Devices to Obtain High-Quality Small Angle X-Ray Scattering (SAXS) Data. Crystals, 10, 975.

58. Putnam, C.D., Hammel, M., Hura, G.L. and Tainer, J.A. (2007) X-ray solution scattering (SAXS) combined with crystallography and computation: defining accurate macromolecular structures, conformations and assemblies in solution. Q Rev Biophys, 40, 191–285.

59. Grant, T.D., Luft, J.R., Carter, L.G., Matsui, T., Weiss, T.M., Martel, A. and Snell, E.H. (2015) The accurate assessment of small-angle X-ray scattering data. Acta Crystallogr D Biol Crystallogr, 71, 45–56.

60. Cantara, W.A., Olson, E.D. and Musier-Forsyth, K. (2017) Analysis of RNA structure using small-angle X-ray scattering. Methods, 113, 46–55.

61. Rambo, R.P. and Tainer, J.A. (2011) Characterizing flexible and intrinsically unstructured biological macromolecules by SAS using the Porod-Debye law. Biopolymers, 95, 559–571.

62. Kikhney, A.G. and Svergun, D.I. (2015) A practical guide to small angle X-ray scattering (SAXS) of flexible and intrinsically disordered proteins. FEBS Lett, 589, 2570–2577.

63. Kozin, M.B., Svergun, D.I. and Embl, B. (2001) Automated matching of high- and low-resolution structural models. J. Appl. Cryst, 34, 33–41.

64. Jones, C.P., Cantara, W.A., Olson, E.D. and Musier-Forsyth, K. (2014) Small-angle X-ray scattering-derived structure of the HIV-1 5’ UTR reveals 3D tRNA mimicry. Proceedings of the National Academy of Sciences of the United States of America, 111, 3395–3400.

65. Spokoini-Stern, R., Stamov, D., Jessel, H., Aharoni, L., Haschke, H., Giron, J., Unger, R., Segal, E., Abu-Horowitz, A. and Bachelet, I. (2020) Visualizing the structure and motion of the long non-coding RNA HOTAIR. Rna, 26, 629–636.

66. Chillón, I. and Marcia, M. (2020) The molecular structure of long non-coding RNAs: emerging patterns and functional implications. Critical Reviews in Biochemistry and Molecular Biology, 55, 662–690.

67. Svergun, D.I. and Nierhaus, K.H. (2000) A Map of Protein-rRNA Distribution in the 70 SEscherichia coli Ribosome*. Journal of Biological Chemistry, 275, 14432–14439.

68. Wang, S., Chan, K.W.K., Tan, M.J.A., Flory, C., Luo, D., Lescar, J., Forwood, J.K. and Vasudevan, S.G. (2022) A conserved arginine in NS5 binds genomic 3’ stem-loop RNA for primer-independent initiation of flavivirus RNA replication. Rna, 28, 177–193.

69. Lee, E., Bujalowski, P.J., Teramoto, T., Gottipati, K., Scott, S.D., Padmanabhan, R. and Choi, K.H. (2021) Structures of flavivirus RNA promoters suggest two binding modes with NS5 polymerase. Nat Commun, 12, 2530.

70. Fajardo, T., Jr., Sanford, T.J., Mears, H.V., Jasper, A., Storrie, S., Mansur, D.S. and Sweeney, T.R. (2020) The flavivirus polymerase NS5 regulates translation of viral genomic RNA. Nucleic acids research, 48, 5081–5093.

