## Supplementary Figures for "Investigating RNA-RNA interactions through computational and biophysical analysis"

**Supplementary Information**

Figure S1. Multiple sequence alignment for 5’TR and 3’TR of 20 mosquito-borne Flaviviruses that was used to form the consensus structure in Fig. 2. The alignment was obtained from the structural alignment toll LocARNA, the two fragments were joined by a linker of 5 Ns. Coloring indicates covariation in consensus base pairs following the RNAalifold schema, ranging from red(no covariation, full primary sequence conservation) to violet (full covariation, all six possible combinations of base pairs). Red boxes highlight the conserved 5’ and 3’ cyclization sequence, respectively.

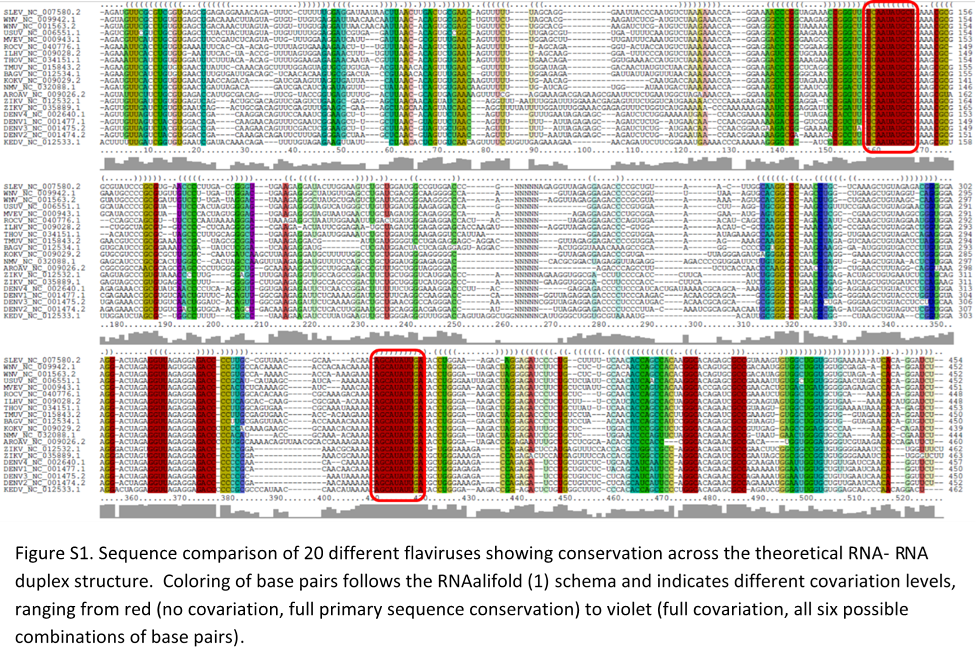

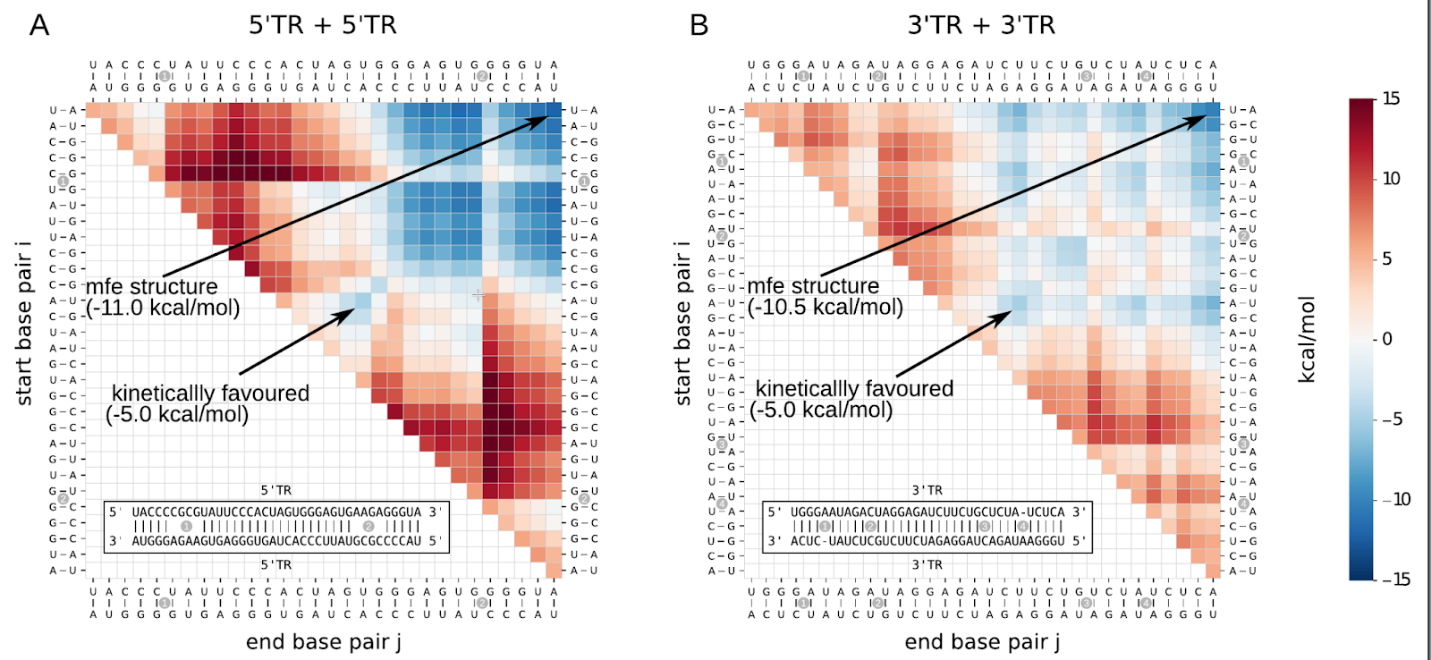

Figure S2. Energy landscapes of predicted homodimer interactions. a) 5’TR homodimer b) 3’TR homodimer. The thermodynamical model predicts stable minimum free energy structures with -11 kcal/mol and -10.5 kcal/mol, for the two respective homodimers. Based on the energy landscapes, there is no kinetically favourable path from a first base pair (drawn on the diagonal) to the full interaction (drawn in the upper right corner) through which the full interaction could be reached without crossing any energy barriers. In both examples, the most stable substructure that is energetically favourable has a stability of -5 kcal/mol and therefore is less stable than the known CS

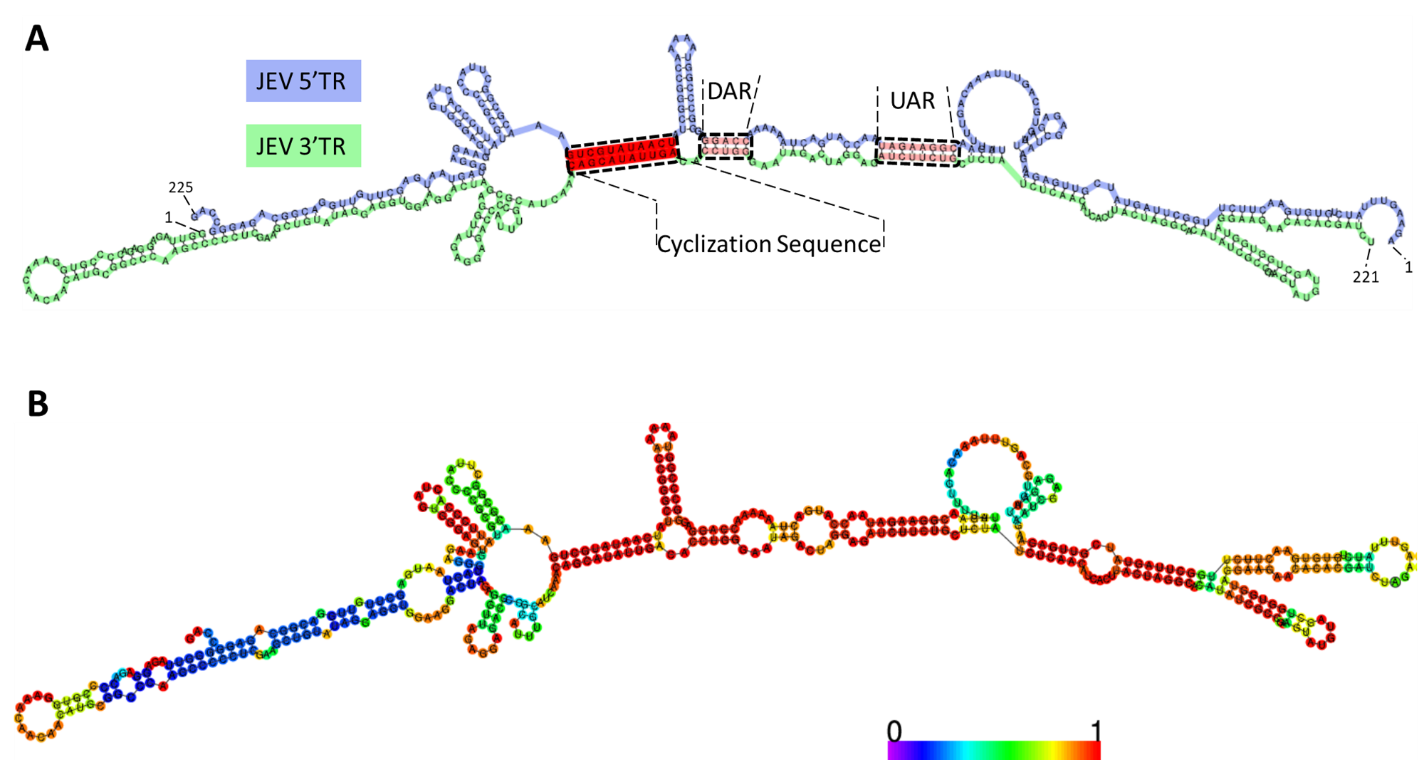

Figure S3. Predicted duplex structure of the JEV 3’TR and JEV 5’TR obtained from RNAcofold, showing the maximum possible extent of interaction. A) Known long range interaction elements, such as cyclization sequence (CS), downstream AUG region (DAR) and upstream AUG region (UAR) are highlighted; B) structure with reliability annotation (based on pair probabilities) ranging from blue (poor reliability) to red (high confidence).

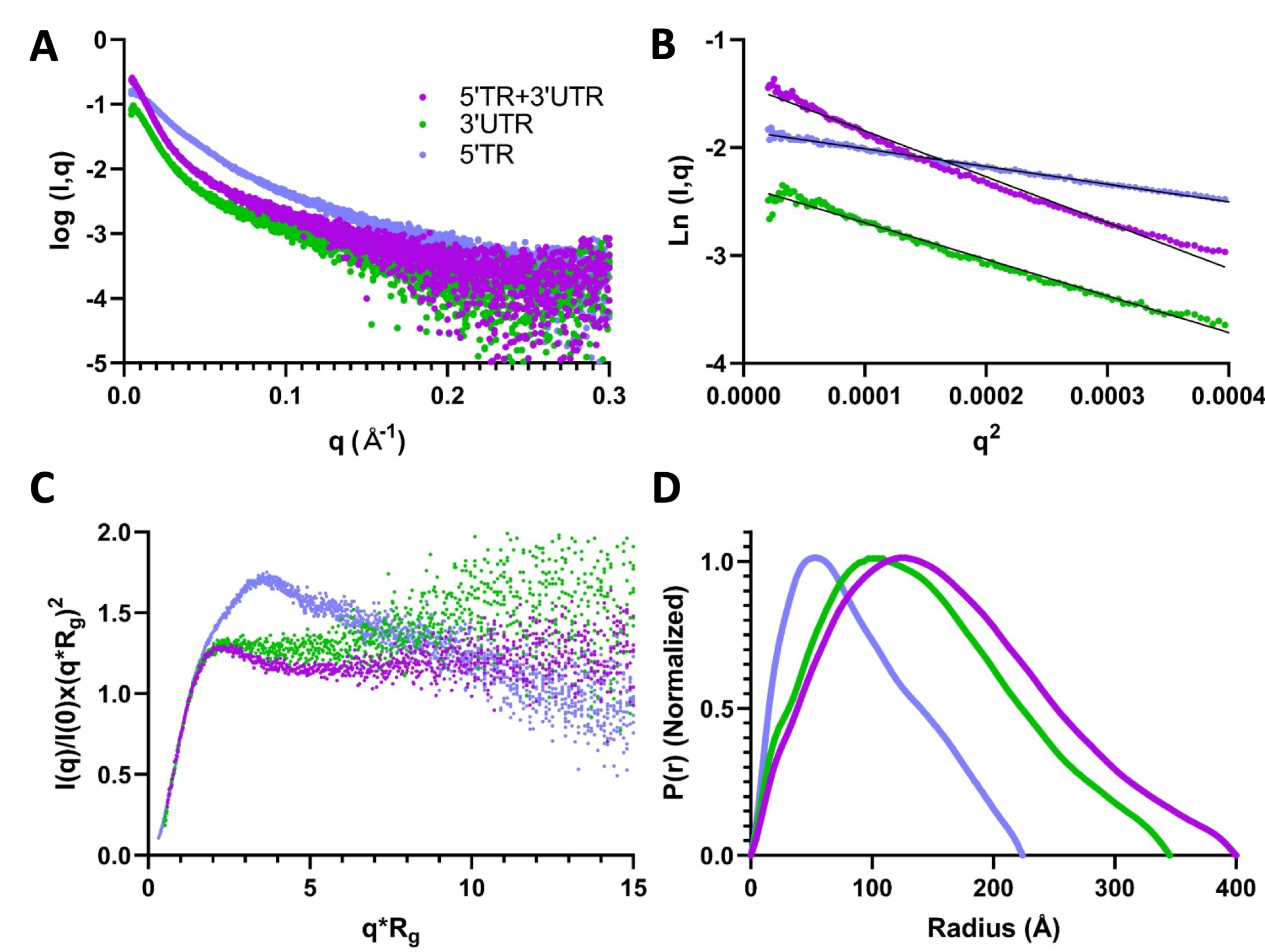

Figure S4. Small Angle X-Ray Scattering (SAXS) plots of JEV RNA-RNA interaction. A) Subtracted & merged scattering data for JEV RNA depicting the scattering intensity vs. scattering angle (q = 4πsinϴ/λ). B) Guinier plots representing the determination of R_g_ and homogeneity derived from the low-angle region. C) Dimensionless Kratky plots of JEV RNA depicting an elongated structure as a result of the non-Gaussian shape(s) of the curve(s). D) Pair-distance distribution plots for JEV RNA for real-space R_g_ and maximal particle dimension (D_max_) determination from the entire SAXS dataset.
